# Xylan modifications in *Populus* that lead to increased growth

**DOI:** 10.1101/2025.03.31.646328

**Authors:** János Urbancsok, Evgeniy N. Donev, Marta Derba-Maceluch, Pramod Sivan, Félix R. Barbut, Madhusree Mitra, Zakiya Yassin, Kateřina Cermanová, Jan Šimura, Michal Karady, Gerhard Scheepers, Ewa J. Mellerowicz

## Abstract

Xylem cells are surrounded by primary and secondary cell walls. Formation of primary walls is regulated by the cell wall integrity surveyance system, but it is unclear if the deposition of secondary walls is similarly regulated. To study this question, we introduced to aspen three different enzymes cleaving cell wall-localized xylan and we suppressed xylan synthase components either ubiquitously or specifically during secondary wall formation. When xylan was ubiquitously altered, 95% of lines showed reduced growth, whereas when it was altered during secondary wall deposition, 30% of lines grew better with the rest having no growth impairment, suggesting opposite effects of primary and secondary wall disturbances. To detect mechanism of growth stimulation by disturbed deposition of secondary wall, we analyzed changes in wood quality traits, chemistry, transcriptomics, metabolomics and hormonomics in transgenic lines. We found increased tension wood production, reduced S- and H-lignin, and changes in several metabolites in common in these lines. Remorin *REM1.3* and *NRL2* (*NPH3* family) transcripts increased and changes in jasmonates, ABA and SA occurred in secondary wall-forming xylem suggesting their involvement in secondary wall integrity surveyance and signaling. The data indicate that a unique program mediates responses to secondary wall impairment that induces growth.

**Highlight:** Disturbance of the deposition of secondary walls that are responsible for the mechanical strength of plant bodies can increase growth with discrete changes in gene expression, hormone levels and metabolism.

## Introduction

Plant cell wall has many essential functions such as protecting protoplasts from abiotic and biotic stresses, regulating cell growth, generating cell shape, providing means for regulation of cellular adhesion, and offering the medium for water and nutrient transport in different tissues. Therefore, cell wall status is expected to be tightly monitored.

Plant cell wall monitoring systems using chemical and mechanical cues have been reported in different types of growing cells (Vaahtera et al., 2019). The best-established mechanism of cell wall status surveillance operates via plasma membrane-located receptor like kinases from the *Catharanthus roseus* receptor-like kinase 1-like family (*Cr*RLK1L) such as THESEUS1 (THE1) and FERONIA (FER). These receptors form signaling complexes interacting among themselves and with other proteins including GPI-anchored co-receptors or LEUCINE-RICH REPEAT EXTENSINs (LRXs) (Doblas et al., 2018). They are thought to perceive wall damage by their extracellular domains interacting with de-methylesterified pectin and anionic peptides called rapid alkalinization factors (RALFs), and to transduce the signals via phosphorylation relay cascades and G-protein-mediated signaling, triggering changes in Ca^2+^fluxes, ROS production in the apoplast and apoplast alkalinization. In the growing pollen tubes and root hairs the integrity of cell wall is directly monitored in the apoplast by RALFs, which bind with high affinity to LRXs and de-esterified pectin and with low affinity to *Cr*RLK1L-GPI-protein signaling complexes providing means for monitoring LRX-homogalacturonan status in the cell wall (Moussu et al., 2023; Schoenaers et al., 2024).

Another well-studied group of proteins perceiving signals from cell wall are different types of wall-associated kinases (WAKs) which are thought to monitor pectin status in the cell wall by interacting with homogalacturonan and its fragments having specific methylesterification status and then transducing signal to the kinase domain in the cytoplasm (Kohorn, 2016a).

Rice WAK11 was shown to regulate the diurnal pattern of growth by controlling activity of brassinosteriod receptor BRI1 in a pectin methylesterification-dependent manner (Yue et al., 2022). Many other chemical signals such as cell wall fragments, peptides or extracellular ATP are generated during growth and pathogen attack. These signals are known as damage-associated molecular patterns (DAMPs) and their perception *via* various pattern recognition receptors has been described in pathways related to cell wall integrity and pathogen sensing which are overlapping and regulate each other (Engelsdorf et al., 2018). The perception of mechanical signals during cell wall integrity stress is less well understood but it likely involves stretch-activated channels such as Ca^2+^-influx MID1-COMPLEMENTING ACTIVITY 1 (MCA1) and REDUCED HYPEROSMOLALITY-INDUCED [Ca^2+^] INCREASE 1 (OSCA1) located in the plasma membrane, which sense low and high osmotic stress, respectively, in the apoplast and activate Ca^2+^-dependent and abscisic acid (ABA) signaling (Vaahtera et al., 2019). Thus, mechanical signaling during cell wall integrity sensing is expected to interact with abiotic stress perception mediated by ABA such as drought, osmotic or freezing stress.

Much less is known about perception of cell wall integrity in non-growing cells such as secondary cell wall-depositing xylem cells. The evidence for such signaling mechanism includes: 1) activation of immune responses and biotic resistance when secondary cell wall is specifically affected (Hernández-Blanco et al., 2007; Gallego-Girlado et al., 2011a, Pogorelko et al., 2013; Pawar et al., 2016; Molina et al., 2021); 2) activation of abiotic stress responses and increased abiotic stress resistance when secondary walls are affected (Chen et al., 2005; Ramírez and Pauly, 2019; Barbut et al., 2024); 3) stimulation of growth or induction of developmental changes in primary-walled cells by alterations of secondary walls (Bosca et al., 2006; Biswal et al., 2015; Derba-Maceluch et al., 2015; Ratke et al., 2018); 4) induction of immune responses (Mélida et al., 2020; Devangan et al., 2023) or developmental changes (Kákošová et al., 2013; Zhao et al., 2013) by secondary wall-derived oligosaccharides and 5) recovery of growth defects of secondary-wall mutants displaying dwarfism by eliminating certain signaling/regulatory proteins and hormones (Gallego-Girlado et al., 2011b; Bonawitz et al., 2014; Ramírez et al., 2018; Ramírez and Pauly, 2019).

In case of lignin mutants that are dwarf, the mechanism of immune responses has been partly elucidated. It involves FER signaling that induces ARABIDOPSIS DEHISCENCE ZONE POLYGALACTURONASE 1 (ADPG1) and ADPG1-mediated release of oligogalacturonides that activate immune defenses in WAK-dependent way (Gallego-Giraldo et al., 2020; Liu et al., 2023). Given the reports of secondary wall xylan modifications leading to increased growth (Biswal et al., 2015; Derba-Maceluch et al., 2015; Ratke et al., 2018) we aimed to investigate mechanisms potentially responsible for this effect. Therefore, we induced different types of xylan alterations in secondary cell walls in aspen and classified them according to their effect on growth. We subsequently selected those transgenic modifications that resulted in growth stimulation and characterized the changes in cell walls, metabolomes, hormonomes and transcriptomes of wood-forming tissues both at the primary and secondary wall stages in the transgenic plants. We detected some common reactions in these plants that could be part of a shared response linking secondary wall defects with increased growth.

## Materials and Methods

### Transgenic lines and wild type

Hybrid aspen (*Populus tremula* L. x *tremuloides* Michx.), clone T89, was used as a wild type. Transgenic lines expressing different fungal xylan-active enzymes (**Table 1**) were generated using *Agrobacterium*-mediated transformation and vectors described in previous publications (Latha Gandla et al., 2015; Donev et al., 2021; Derba-Maceluch et al., 2023; Barbut et al., 2025; Sivan et al., 2025). Lines with the highest transgene expression or native gene suppression were selected from at least 20 independent lines per construct. Lines with suppressed native xylan biosynthetic *GT43* genes were selected among the lines described in an earlier publication (Ratke et al., 2018).

**Table 1.**
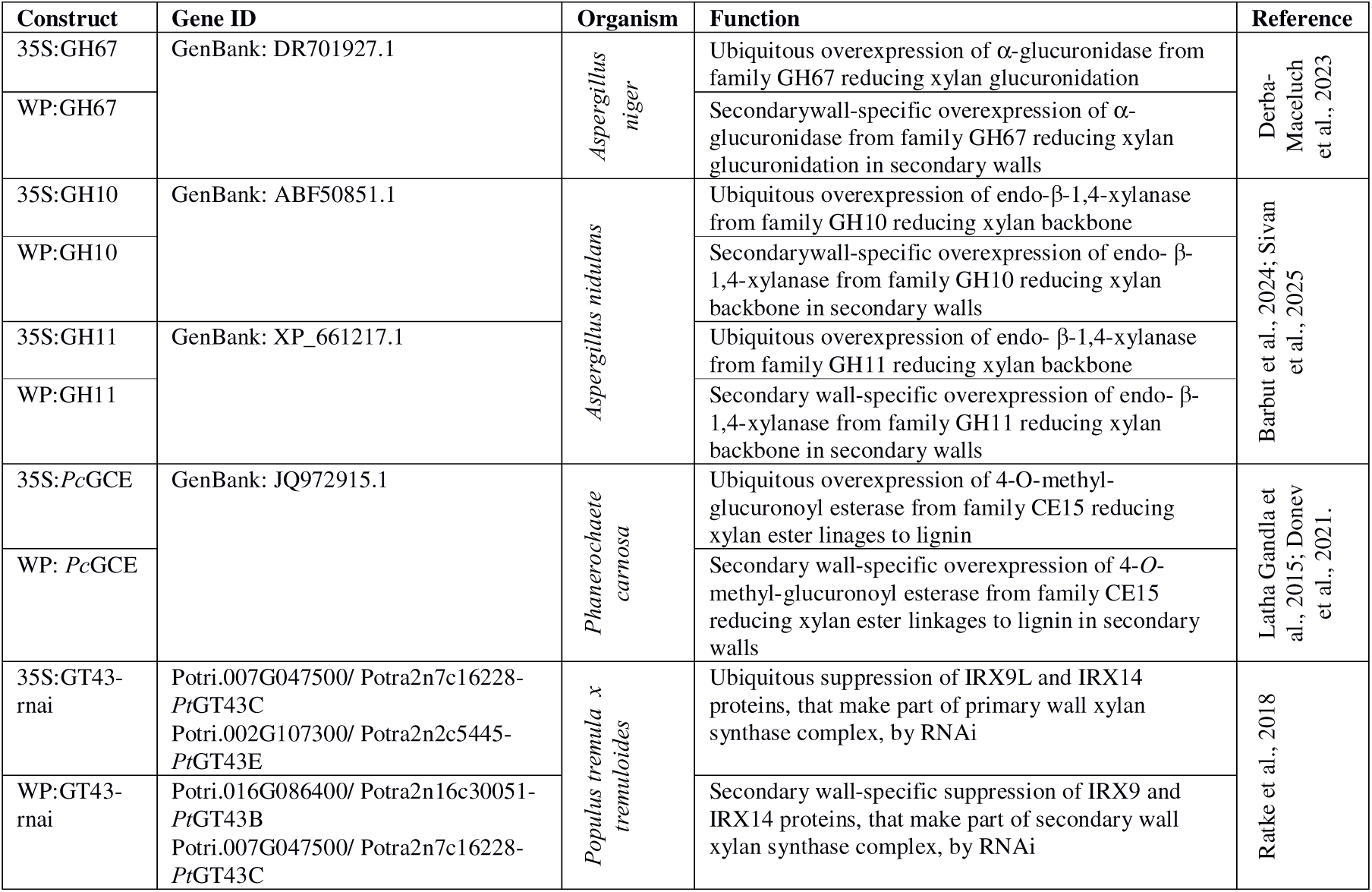
List of constructs used in this study.

### Greenhouse conditions, growth analyses and tissue collection

*In vitro* propagated saplings were planted in soil, and grown for nine weeks in the phenotyping platform (WIWAM Conveyor, custom designed by SMO, Eeklo, Belgium) as described by Wang et al. (2022) and Urbancsok et al., (2023). Briefly, the conditions were 18 h /6 h (day/night) with light having 160-230 µmol m^-2^ s ^-1^ intensity during the day provided by white light (FL300 LED Sunlight v1.1) and far-red light (FL100 LED custom-made, 725-735 nm) from Senmatic A/S (Søndersø, Denmark), 22 °C /18 °C temperature, and the average air relative humidity of 60%. Plants were watered automatically based on weight and their height was automatically measured.

At harvest, the trees were photographed, and their stem diameters (at the base) and aboveground fresh weights were recorded. For RNA, hormonomics and metabolomics analyses, a 30 cm-long stem segment above internode 37 was dissected, debarked, frozen in liquid nitrogen and stored at -70 °C. Frozen bark and frozen wood core were scraped with a scalpel into a precooled mortar and ground in liquid nitrogen to obtain frozen cambium/phloem and developing xylem powder, respectively.

The stem below internode 37 was used for determining average internode length. For SilviScan analysis, the four-cm long segment from the base of stem was used, and the remaining stem was debarked and freeze-dried for 48 h for wood chemistry analyses. Belowground biomass was determined by weighing cleaned and air-dried roots.

### RT-PCR

To isolate total RNA, frozen cambium/phloem and xylem powders (approximately 100 mg) were extracted with CTAB/chloroform:isoamylalcohol (24:1), followed by precipitation with LiCl and sodium acetate/ethanol (Chang et al., 1993). RT-PCR procedures followed protocol described by Sivan et al., (2025). UBQ-L (Potri.005G198700) and ACT11 (Potri.006G192700) were selected as reference genes. The primer sequences are listed in **Supplementary Table S1**. The relative expression level was calculated according to Pfaffl (2001).

### SilviScan and NIR analysis

Wood quality traits were analyzed using a SilviScan instrument (RISE, Stockholm, Sweden) as described by Urbancsok et al. (2023) for six transgenic trees per line and 12 wild type trees. The same samples were scanned by a near infrared (NIR, 960-2500 nm at 256 wavelengths) scanner at 30 µm resolution and tension wood was predicted for each pixels using a procedure described by Renström et al, 2025. Heatmaps showing tension wood distributions were subsequently produced and average probability of tension wood was calculated for each sample.

### Cell wall chemical analyses

Wood was analyzed in three trees per line, as described by Sivan et al. (2025). Briefly, the wood powder was obtained by filing the freeze-dried wood and sieving the sawdust with Retsch AS 200 analytical sieve shaker (Retsch GmbH, Haan, Germany) to the particle size between 50 and 100 µm. Approximately 50 µg (± 10 µg) of this powder was used in a pyrolizer (PY-2020iD and AS-1020E, Frontier Lab, Japan) connected to a GC/MS (7890A/5975C, Agilent Technologies Inc., Santa Clara, CA, USA) to analyze main wood components including carbohydrates, H-, G- and S-lignin units, and phenolic compounds, as described by Gerber et al. (2012). The composition of matrix polysaccharides was analyzed using the methanolysis-trimethylsilyl (TMS) procedure as described by Pramod et al. (2021). Briefly, the alcohol-insoluble residue (AIR) prepared according to Latha Gandla et al. (2015) was destarched by α-amylase (from pig pancreas, cat. nr. 10102814001, Roche, USA) and amyloglucosidase (from *A. niger* cat. nr.10102857001, Roche) enzymes, and used to prepare silylated monosaccharides that were separated by GC-MS (7890A/5975C; Agilent Technologies Inc., Santa Clara, CA, USA) as described in Latha Gandla et al. (2015). Raw data MS files from GC-MS analysis were converted to CDF format in Agilent Chemstation Data Analysis (v.E.02.00.493) and exported to R software (v.3.0.2). 4-*O*-methylglucuronic acid was identified according to Chong et al. (2013).

### Hormonomics

Frozen cambium-phloem and developing xylem tissue powders were extracted and subjected to GC-MS analysis as described by Šimura et al. (2018), with slight modifications (Urbancsok et al., 2023). The plant hormone ethylene’s immediate non-volatile precursor, 1-aminocyclopropane-1-carboxylic acid (ACC), was measured using liquid chromatography-tandem mass spectrometry (LC-MS/MS), following the methodology in Karady et al. (2024). All identified hormones including their precursors and inactivated forms are listed in **Supplementary Table S2**.

### Metabolomics

Metabolic profiling was performed by GC-MS and LC-MS at the Swedish Metabolomics Centre, Umeå, Sweden as described previously (Gullberg et al. 2004). Approximately 10 mg of cambium/phloem and developing xylem tissue was extracted using an extraction buffer (20:20:60, chloroform:water:methanol, v/v/v) with internal standards. For LC-MS, the samples were analyzed using an Agilent 1290 Infinity UHPLC system (Agilent Technologies, Waldbronn, Germany) coupled to a 6546 Q-TOF mass spectrometer (Agilent Technologies, Santa Clara, CA, USA), with chromatographic and mass spectrometric parameters optimized for both positive and negative ion modes. GC-MS analysis involved derivatized samples analyzed on an Agilent 7890B gas chromatograph (Restek Corporation, Bellefonte, PA, USA) coupled to a Pegasus BT TOF-MS (Leco Corp., St Joseph, MI, USA), with detailed instrumental settings and data processing performed using ChromaTOF, MATLAB (Mathworks, Natick, MA, USA), and NIST MS 2.2 software (https://chemdata.nist.gov/mass-spc/ms-search/downloads/)

Data processing for LC-MS included targeted feature extraction using in-house libraries, while GC-MS employed retention index and mass spectral comparisons for metabolite identification as described in Schauer et al. (2005). Further technical details, including specific instrument settings and data processing methodologies, are provided in the supplementary document.

### Transcriptomics

For transcriptomics, RNA from five trees per transgenic line and eight trees from wild type was purified as described by Urbancsok et al. (2023) and used for cDNA preparation and sequencing at Novogene Co., Ltd. (Cambridge, United Kingdom). Quality control and mapping to the *P. tremula* transcriptome (v.2.2), retrieved from the PlantGenIE (https://plantgenie.org; Sundell et al., 2015) and to sequences used in fungal vectors were carried out by Novogene. Raw counts were used for differential expression analysis in R (v.3.4.0) using Bioconductor (v.3.4) DESeq2 package (v.1.16.1), as described by Kumar et al., 2019).The best BLAST hits were identified in *Populus trichocarpa* (v.3.1) and *Arabidopsis thaliana* (v.11.0).

A weighted gene correlation network analysis (WGCNA) was obtained using R [v.3.4.0; https://www.R-project.org] library WGCNA (Langfelder & Horvath, 2008). Gene modules were correlated with traits from growth, hormonomics, metabolomics and SilviScan data. Genes from selected modules were used to design co-expression gene networks obtained using PlantGenIE tools. The networks were visualized by Cytoscape (v.3.6.0) (Shannon et al., 2003).

### Statistical analyses

Unless otherwise stated, univariate statistical analyses were performed in JMP Pro (v.16.0) software (SAS Institute Inc., Cary, NC, USA).

## Results

### Growth of transgenic lines with modified xylan

To investigate which types of xylan modification increase growth, we used 43 transgenic hybrid aspen lines carrying ten different constructs that were designed to induce different types xylan modifications (**Table 1**). Majority of these lines were expressing different fungal enzymes acting on xylan and targeted to cell walls using the previously described strategies (Derba-Maceluch et al., 2023; Barbut et al., 2024; Sivan et al., 2025). In other lines, different members of xylan synthase complex from family GT43 were suppressed (Ratke et al., 2018). The transgenes were expressed by either a ubiquitous (35S) promoter or the wood-specific (WP) promoter active during secondary wall biosynthesis (Ratke et al., 2015). In case of native GT43 genes, the 35S construct targeted the primary wall xylan synthase genes, *PtGT43C* and *E*, whereas the WP construct affected expression of the secondary wall xylan synthase genes *PtGT43B* and *C* (Ratke et al., 2018).

The modification of xylan differentially affected growth, depending on the kind of transgene, the type of promoter and the level of transgene expression (**Figure 1a, b**). *GH67* transgene encoding xylan α-glucuronidase induced growth when highly expressed specifically in the secondary walled tissues, but not when 35S promoter was used. The transgene expression level was similar or higher in 35S promoter-driven lines compared to WP. Endo-β-1,4-xylanases (GH10 and GH11) by large inhibited growth regardless the promoter, and their effect was proportional to transgene expression. The exception was a low-expressing GH10 line with WP, which had significantly increased height compared to wild-type. Glucuronoyl esterase-expressing lines exhibited premature senescence when 35S promoter was used as was previously observed (Latha Gandla et al., 2015; Donev et al., 2021) and their height was affected in one of the studied lines. Interestingly, the same transgene driven by WP showed growth stimulation (**Figure 1a**). In this case, the phenotype was dependent on the choice of promoter rather than the transgene expression level. The suppression of primary wall xylan synthase complex members, *Pt*GT43C and *Pt*GT43E, using 35S promoter RNAi constructs resulted in growth inhibition or no effect, but the suppression of the members of secondary wall xylan synthase complex, *Pt*GT43B and *Pt*GT43C, using WP led to enhanced growth as previously observed (Ratke et al, 2018). In summary, for four out of five transgenes there was a significant difference between promoters, with better growth when WP was used, and all the best performing individual lines had WP-driven transgenes. For all four fungal genes studied here, the expression levels were lower when WP was used compared to 35S promoter, whereas *PtGT43C* native gene suppression was not affected by the promoter. Thus, the data support the hypothesis that xylan modification induced specifically in secondary walls can increase growth but this is conditional to transgene type and its expression level.

**Figure 1.**
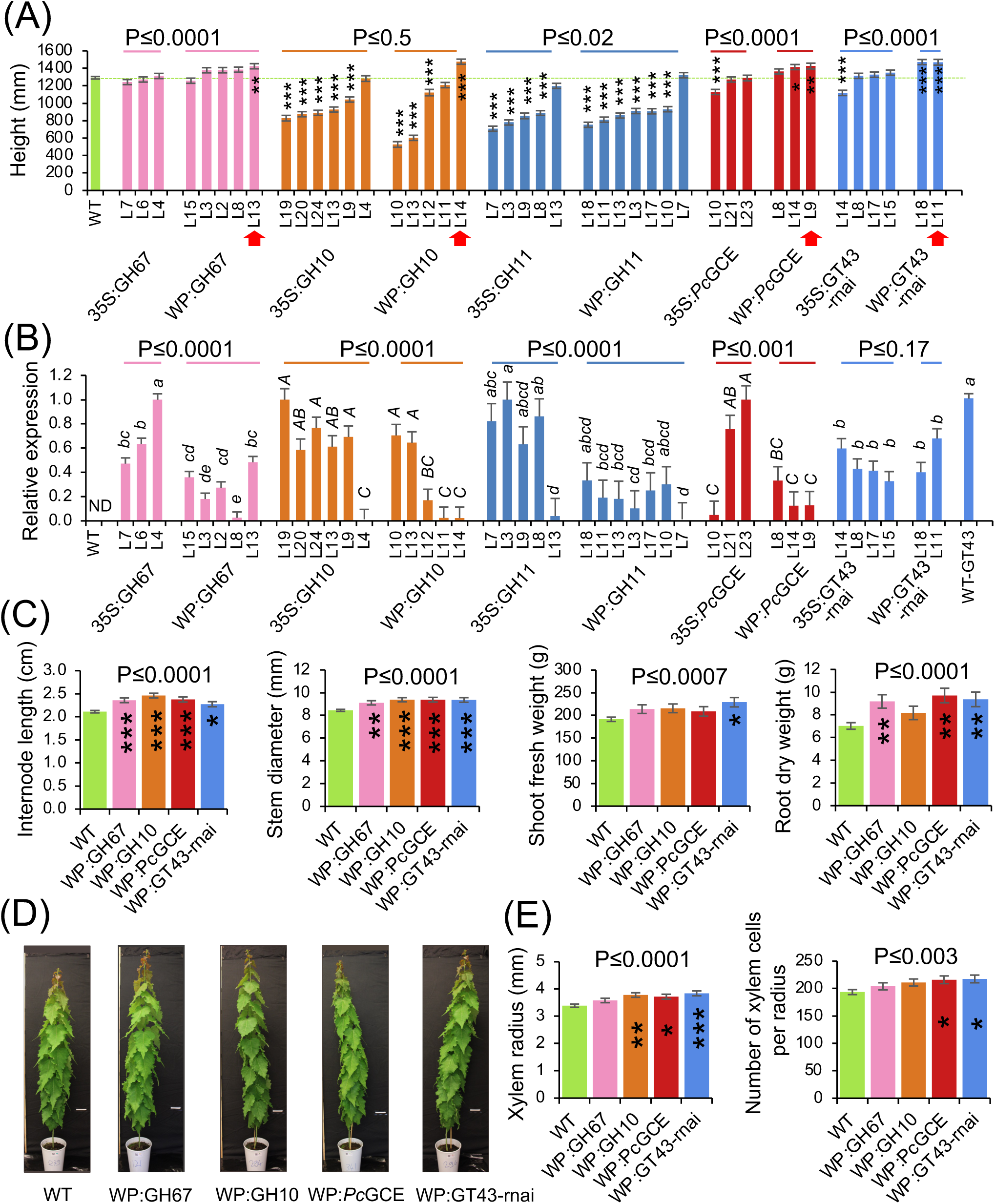
Xylan modification affects growth of hybrid aspen. **(A)** Height of transgenic lines with modified xylan either ectopically (35S) or specifically in secondary walls (WP). Red arrows indicate lines with increased growth selected for further analysis. **(B)** Relative expression levels of transgenes in lines overexpressing fungal enzymes (GH67, GH10, GH11, *Pc*GCE) or targeted native *PtxtGT43C* normalized to the highest-expressing line. ND – transgenes were not detected in WT samples. For each transgene/gene, the means accompanied by the same letter are not significantly different (P≤0.05, Tukey’s test). **(C)** Different morphological parameters of selected lines with increased growth. **(D)** Representative individuals of selected lines. **(E)** Xylem production determined by SilviScan analysis of wood in selected lines. Data are means ± SE; N=6, 3, or 6 trees for transgenic lines and 24, 6 or 12 trees for WT in (A,C), (B), or (E), respectively. * - P≤0.05; ** - P≤0.01; *** - P≤0.001 for comparisons with WT by Dunnett’s test. P values above the bars show significance of differences between 35S and WP promoters (A, B) or between WT and all selected lines (C, E) (post-ANOVA contrast). The height data for some lines expressing xylanases have been reported by Sivan et al. (2025) and are shown here for the comparative purpose.

Four transgenic lines with different types of transgenes all exhibiting increased stem height were subsequently selected for thorough phenotypic analyses. The selected individual lines had increased internode length and stem diameter, and collectively they had greater shoot fresh weight and root dry weight than wild type but showed no alterations in general morphology (**Figure 1c, d**). Wood quality traits analyzed by SilviScan showed no effect of genotype in any of the analyzed parameters indicating that wood anatomy and physical properties did not differ in xylan-modified lines with increased growth from type (**Supplementary Table S3**). However, there was an increase in xylem radius and number of xylem cells per radius in three or two lines, respectively, and when comparing all transgenic lines combined to wild-type plants, indicating increased xylem production by the cambium in transgenic lines with increased height growth (**Figure 1e**).

As young aspen trees typically form some tension wood even when growing upright, we wondered if tension wood formation was affected in transgenic lines with modified secondary wall xylan and increased growth. To test it, we used NIR spectroscopy of the cut stem surface at the base of the stems that allows us to accurately predict the occurrence of tension wood based on cell wall chemical fingerprints (Renström et al., 2025). The heatmaps showing the distribution of probability of tension wood and its quantification indicated a greater occurrence of this tissue in the transgenic lines compared to wild type (**Figure 2**).

**Figure 2.**
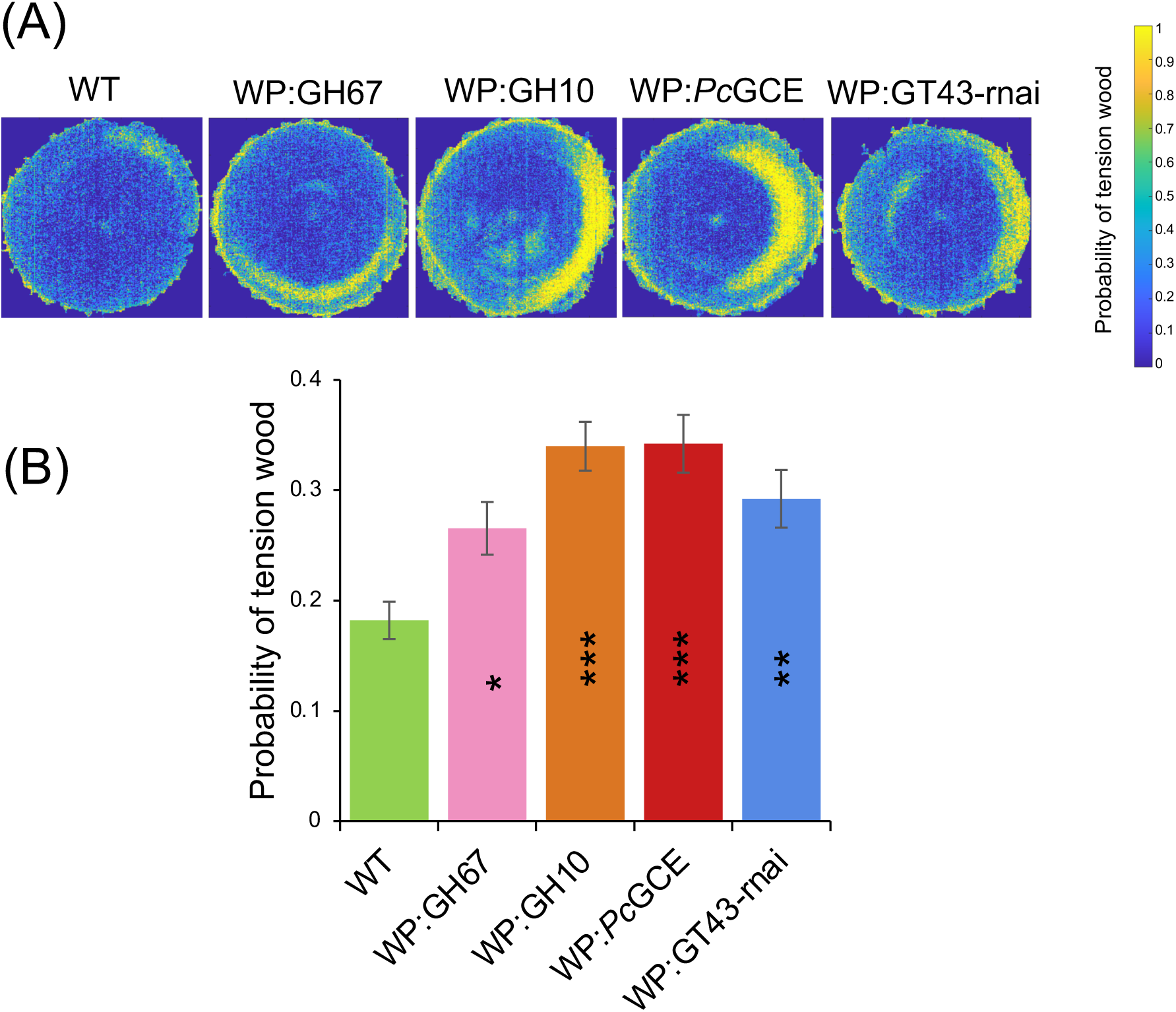
Transgenic lines with xylan defects in secondary walls and increased growth produce more tension wood compared to wild type. **(A)** Tension wood probability shown as heatmaps of representative NIR images for transgenic lines and wild type (WT). **(B)** Quantification of tension wood probability. Data are means ± SE; N=6 trees for transgenic lines and 12 trees for WT. * - P≤0.05; ** - P≤0.01; *** - P≤0.001 for comparisons with WT by Dunnett’s test.

### Wood cell wall composition analyses in xylan-modified lines with increased growth show slightly reduced lignification

The transgenic modifications of selected lines were targeted to developing secondary walls hence we analyzed their wood cell wall composition. Matrix polysaccharides were only slightly affected in GH67-overexpressing line that exhibited an increase in mannose and reductions in 4-*O*-methyl-glucuronic acid (meGlcA) (expected based on the GH67 enzyme activity) and galacturonic acid (GalA) (**Figure 3a**). GH10 induced increases in mannose and pectin-related sugars: arabinose, rhamnose and GalA. *Pc*GCE induced increased glucuronic acid, and suppression of endogenous *GT43B* and *C* genes resulted in a small decrease in xylose as observed before (Ratke et al., 2018). Overall, these changes were small and specific for each transgene.

**Figure 3.**
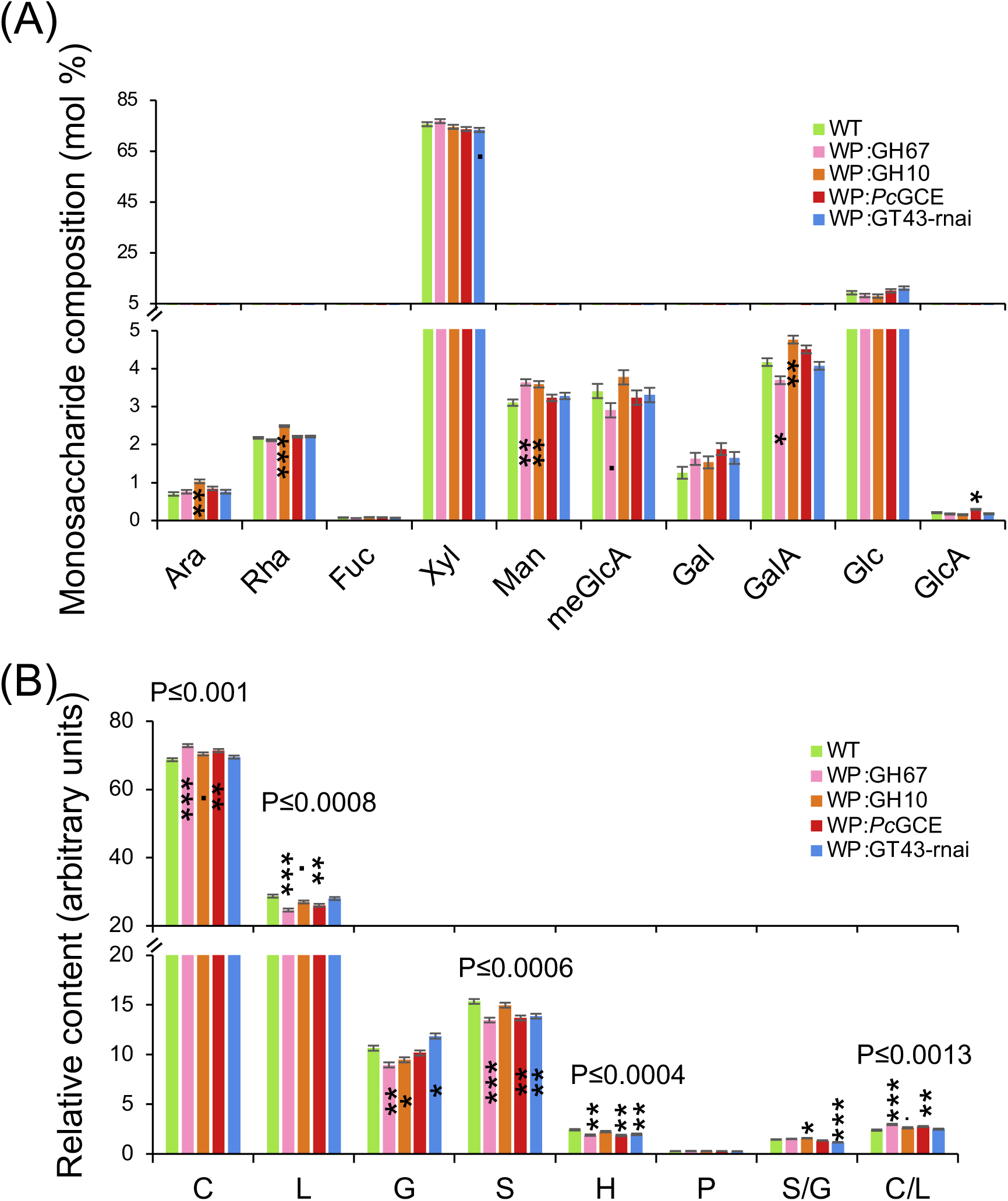
Changes in wood cell wall composition in xylan-modified lines with increased growth. **(A)** Monosaccharide composition of matrix polysaccharides analyzed by methanolysis-trimethylsilyl analysis. **(B)** Cell wall composition according to pyrolysis-GC/MS. Data are means ± SE; N=3. Significant differences for individual lines compared to wild type (WT) according to Dunnett’s test are indicated as follows: · - P≤0.1; - P≤0.05 ; * - P≤0.05; ** - P≤0.01; *** - P≤0.001. P values above the bars in (B) show significance of differences between WT and all lines collectively (post-ANOVA contrast). Ara - arabinose; Fuc - fucose; Gal - galactose; GalA - galacturonic acid; Glc - glucose; GlcA – glucuronic acid; Man - mannose; MeGlcA - methylglucuronic acid; Rha - rhamnose; Xyl - xylose. C- carbohydrates; G- guaiacyl lignin units; H - *para-*hydroxyphenyl lignin units; L - total lignin (S+G+H); P-– phenolics; S - syringyl lignin units.

Despite the transgenes targeting secondary wall xylan in different ways, some common changes in cell wall were revealed by pyrolysis-GC/MS analysis. The overall carbohydrate content of the transgenic lines seemed to be greater than in wild type, which was at the expense of lignin especially the S- and H-lignin units (**Figure 3b**). In contrast, the G-lignin content increased in the *PtGT43*-suppressed line but decreased in GH67- and GH10-expressing lines resulting in variable trends in S/G lignin ratio among the lines.

### Hormonomics analyses in transgenic lines with increased growth indicate changes in cytokinins, jasmonates, ABA and SA

To investigate if different types of xylan modification in secondary walls affected similar hormonal pathways, we carried out hormonomics analyses (Šimura et al., 2018) detecting different forms of cytokinins, auxins, jasmonates, salicylic acid (SA), abscisic acid (ABA) and 1-aminocyclopropane-1-carboxylic acid (ACC) (**Supplementary Table S2**). Separate cambium/phloem samples including cells with primary walls, and developing xylem samples containing cells depositing secondary walls were analyzed. Among the cytokinins, the cytokinin precursor *cis*-zeatin riboside (*c*ZR), which was the most abundant species, increased in two or three out of four tested lines in the cambium/phloem and xylem tissues, respectively, with a similar tendency detected for all analyzed lines in both tissues (**Figure 4**). IAA and its oxidation product oxIAA had a tendency to decrease in the cambium/phloem.

**Figure 4.**
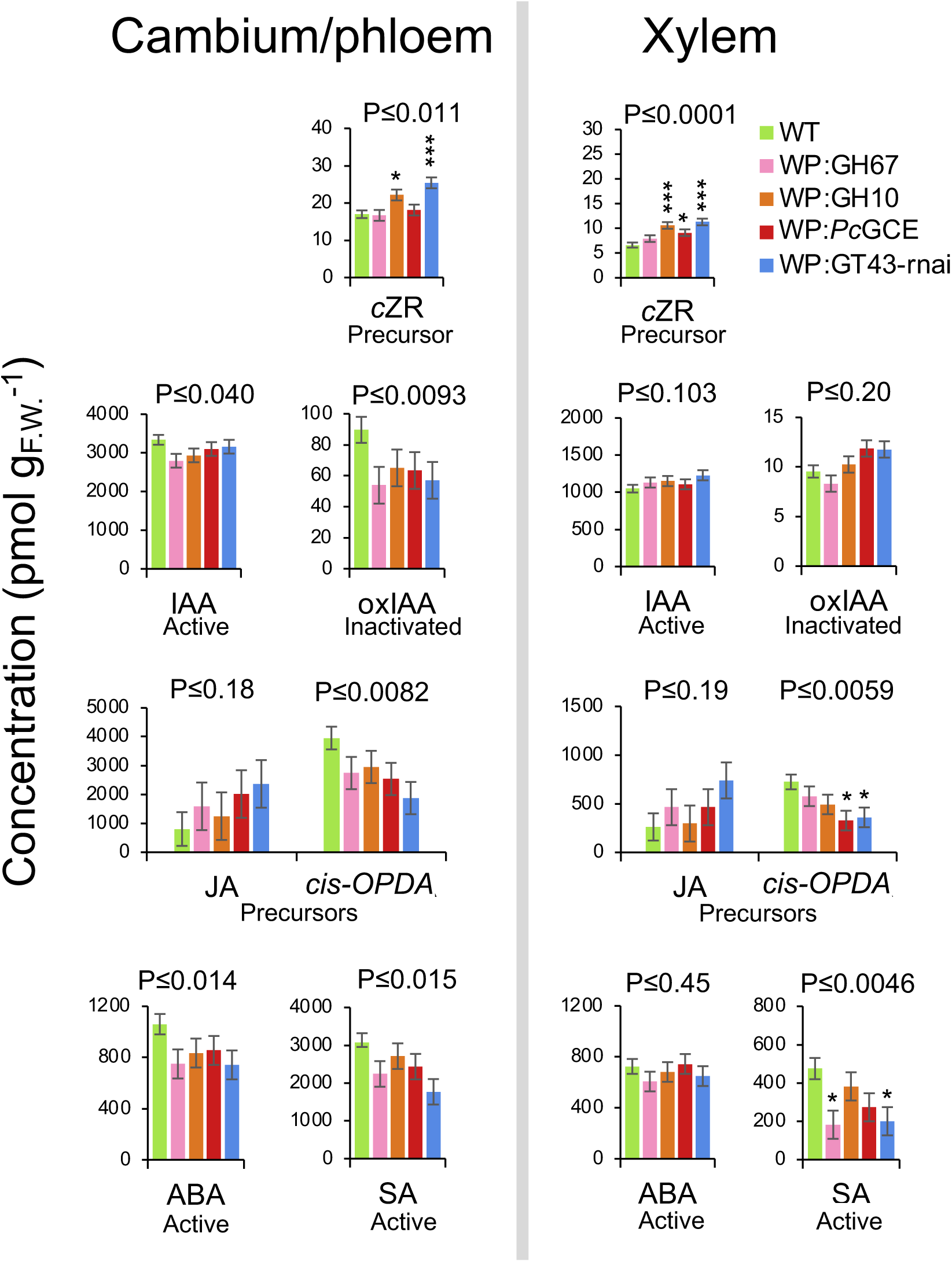
Changes in hormonal status in wood-forming tissues of xylan-modified lines with increased growth. Data are means ± SE; N=4 for transgenic lines and 8 for WT. Asterisks indicate significance of differences compared to wild type (WT) by Dunnett’s test: * - P≤0.05; ** - P≤0.01; *** - P≤0.001. P values above the bars show significance of differences between WT and all lines (post-ANOVA contrast). ABA - abscisic acid; *cis-* OPDA - *cis*-12-oxophytodienoic acid; *c*ZR - *cis*-zeatin riboside; IAA - indole-3-acetic acid; JA - jasmonic acid; oxIAA - oxidized IAA; SA - salicylic acid.

The precursor of jasmonic acid (JA), *cis*-12-oxophytodienoic acid (*cis*-OPDA), decreased significantly in two transgenic lines in the xylem with a similar pattern observed for all lines in both analyzed tissues, whereas its downstream reaction product JA showed some tendency for increase, but the levels of JA were very variable among individual plants resulting in high probability values. SA decreased significantly in two transgenic lines in the xylem and all lines showed similar tendency in both tissues. ABA contents tended to be reduced in the cambium/phloem of transgenic lines compared to wild type, whereas the changes were negligible in the developing xylem tissue. ACC was affected only in the line WP:*Pc*GCE that had higher levels of this precursor than wild type in the xylem (**Supplementary Table S2**). To conclude, the common changes in different xylan-modified lines compared to wild type included elevated *c*ZR contents in both tissues, decreased IAA and oxIAA amounts in the cambium/phloem, reduced levels of *cis*OPDA in both tissues, likely with the biosynthesis of JA, as well as reduced contents of ABA in the cambium/phloem and SA in both tissues.

### Xylan-modified lines with better growth performance exhibited common changes in metabolites

To further investigate whether different types of xylan modification in secondary walls resulting in better growth affected similar metabolic pathways or not, we carried out metabolomics analyses using both gas and liquid chromatography. Among the 300 metabolites identified by these analyses in the wood-forming tissues (cambium/phloem and developing xylem), 64 were affected in transgenic samples specifically in the cambium/phloem tissues, 21 were affected only in the developing xylem and 20 were affected in both tissues when compared to wild type (**Figure 5a**). The metabolites were classified (**Supplementary Table S4**) and the affected groups are shown as volcano plots (**Figure 5b**). In the cambium/phloem tissues, there was a decreased abundance in amino acids and increased phenolic glycosides. Some lignols and other phenylpropanoids were also slightly increased in both tissues. The affected amino acids were those that served as phenylpropanoid precursors. (phenylalanine and threonine). They were altered in the cambium/phloem only whereas methionine was reduced in both tissues. In addition, some monosaccharides increased in abundance in the transgenic lines including xylose, arabinose and arabitol in both tissues, also xylobiose and a related xylitol increased in the cambium/phloem tissues. The observed metabolic alterations point towards common changes in the phenylpropanoid pathway in xylan-impaired lines that led to higher levels of phenolic glycosides, and in xylan-related sugars. Last but not least, the metabolomics analysis confirmed the reduced salicylic acid level in the xylem (**Supplementary Table S4**).

**Figure 5.**
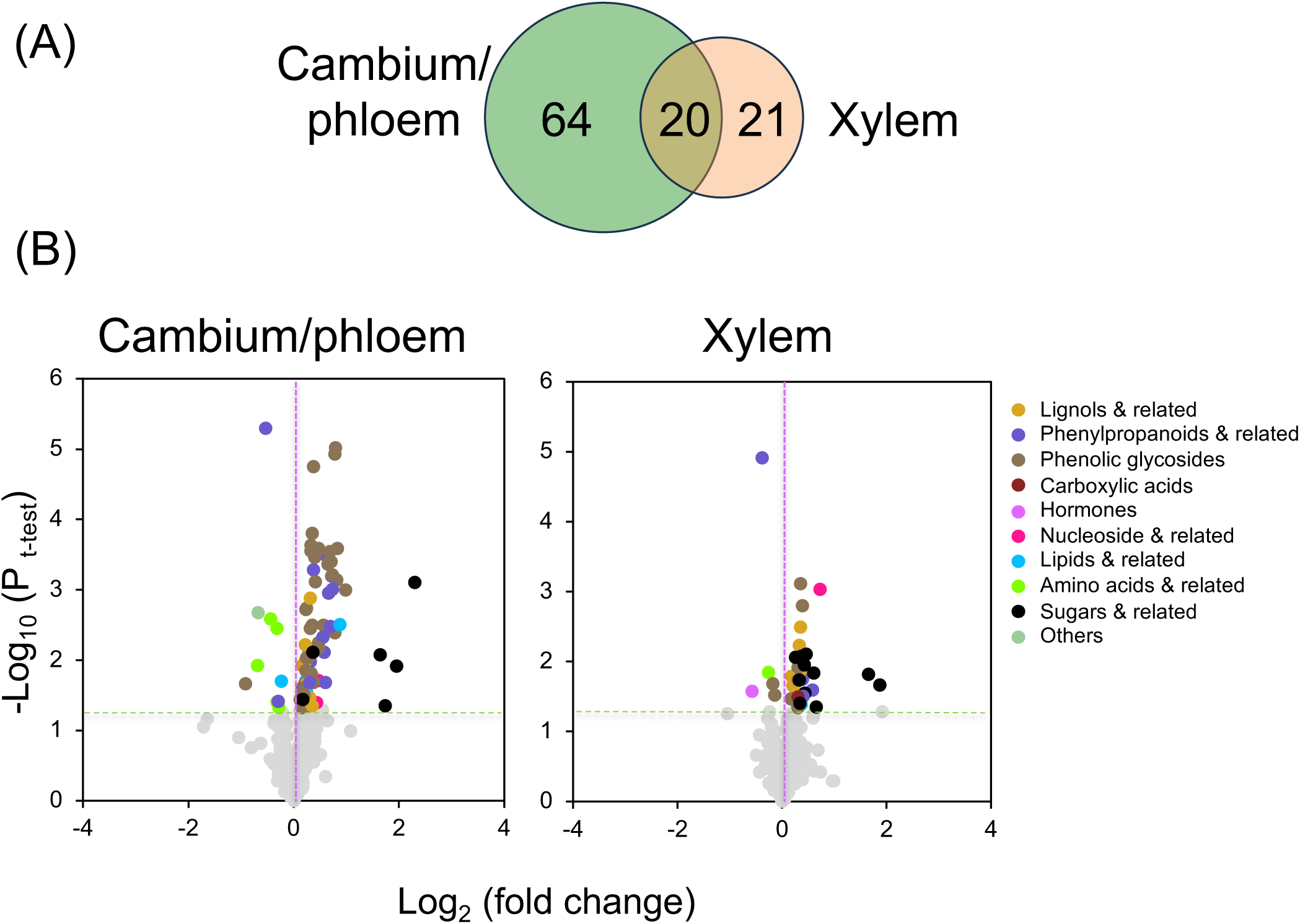
Metabolomes of cambium/phloem and developing xylem in transgenic lines with altered xylan and increased growth show changes in several groups of compounds. **(A)** Venn diagram showing number of metabolites affected in transgenic lines in cambium/phloem and developing xylem tissues out of 300 detected metabolites. **(B)** Volcano plots of metabolites analyzed by LC-MS and GC-MS showing groups of compounds significantly affected (P≤0.05, t-test) in samples of transgenic lines (WP:GH67, WP:GH10, WP:*Pc*GCE, WP:GT43-rnai) taken together compared to wild type. Data are based on 8 WT trees and 4 transgenic trees in each line and tissue. The list of affected compounds is given in **Table S4**.

### Commonly affected transcripts detected in transgenic lines with better growth

To reveal if the different xylan modifications in the transgenic lines with enhanced growth induced common transcriptomic changes, RNA sequencing was performed in the cambium/phloem and developing xylem tissues.

In the cambium/phloem of transgenic xylan-modified, between 6 and 136 genes were differentially expressed compared to wild type (P_adj_ ≤ 0.01, | log_2_(fold change) | ≥ 0.584) and no commonly affected genes were observed (**Figure 6, Supplementary Table S5**). In contrast, in developing xylem, between 17 and 128 genes were differentially expressed, and three genes were affected in common in all the lines. One of them has no assigned function and no homologues in other species but poplars. The second gene is similar to *Arabidopsis NRL2* from the *NPH3* family of plant-specific adapter proteins and the third one is similar to *Arabidopsis Remorin 1.3* (*REM1.3*) regulating lipid raft microdomain organization in the plasma membrane. These three genes are potentially regulated by the secondary cell wall integrity impairment signals *in situ*.

**Figure 6.**
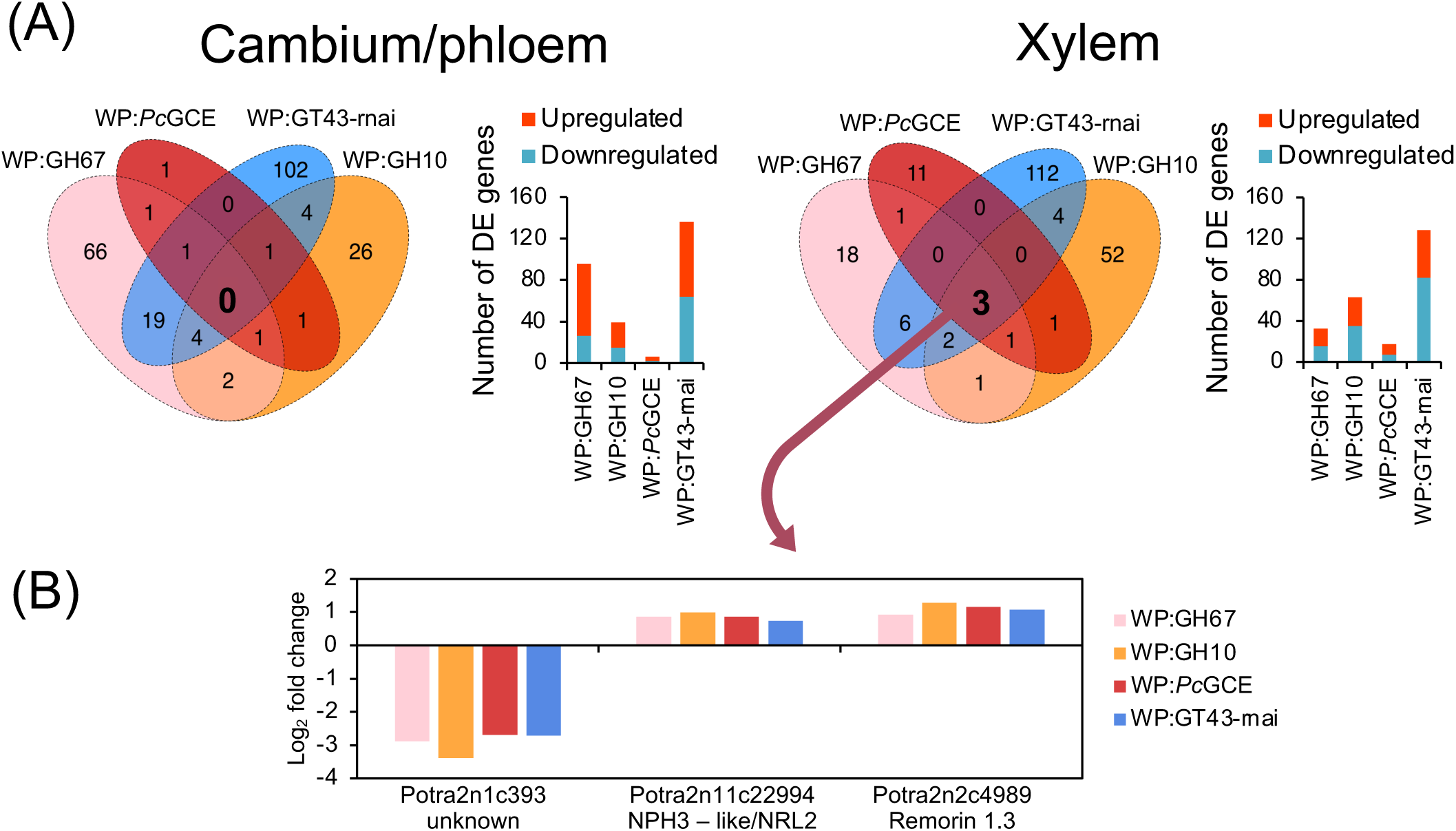
RNA sequencing of wood-forming tissues in xylan-modified lines with increased growth. **(A)** Venn diagrams showing numbers of differentially expressed (DE) genes compared to wild type (WT) in each line (P_adj_ ≤ 0.01,|Log_2_(fold change) ≥ 0.584) and bar charts showing numbers of up- and downregulated genes in each line. **(B)** The differential expression of three genes affected in common in the xylem of transgenic lines.

To further explore transcriptomic changes in the transgenic lines, we carried out a weighted gene correlation network analysis (WGCNA) (Horvath, 2011) in the cambium/phloem and developing xylem tissues. This analysis groups genes with highly correlated expression patterns into color-coded clusters. The expression of each cluster is then summarized using the eigengene values which serve to create dendrograms showing similarities among the samples and to analyze correlations to external sample traits, in our case growth, SilviScan, hormonomics and metabolomics data. The WGCNA clusters for three hierarchical levels are shown in **Figure 7a**. The co-regulated genes for the middle level of clustering are listed in **Supplementary Tables S6 and S7** for the cambium/phloem and xylem tissues, respectively. In the cambium/phloem, 14 clusters were detected (**Figure 7b**) and one of them, the Tan cluster, grouped all transgenic samples separate from the wild-type samples (**Figure 7c, Figure S1**). Among the external variables correlated with the eigengene values of the Tan cluster, there were internode length, phenylalanine, many different phenylpropanoids, and hormones *t*Z and IAA (**Supplementary Table S8**). In the xylem samples, seven gene clusters were detected but no grouping could be found separating transgenic lines from wild type (**Figure 7b**, **Figure S2**).

**Figure 7.**
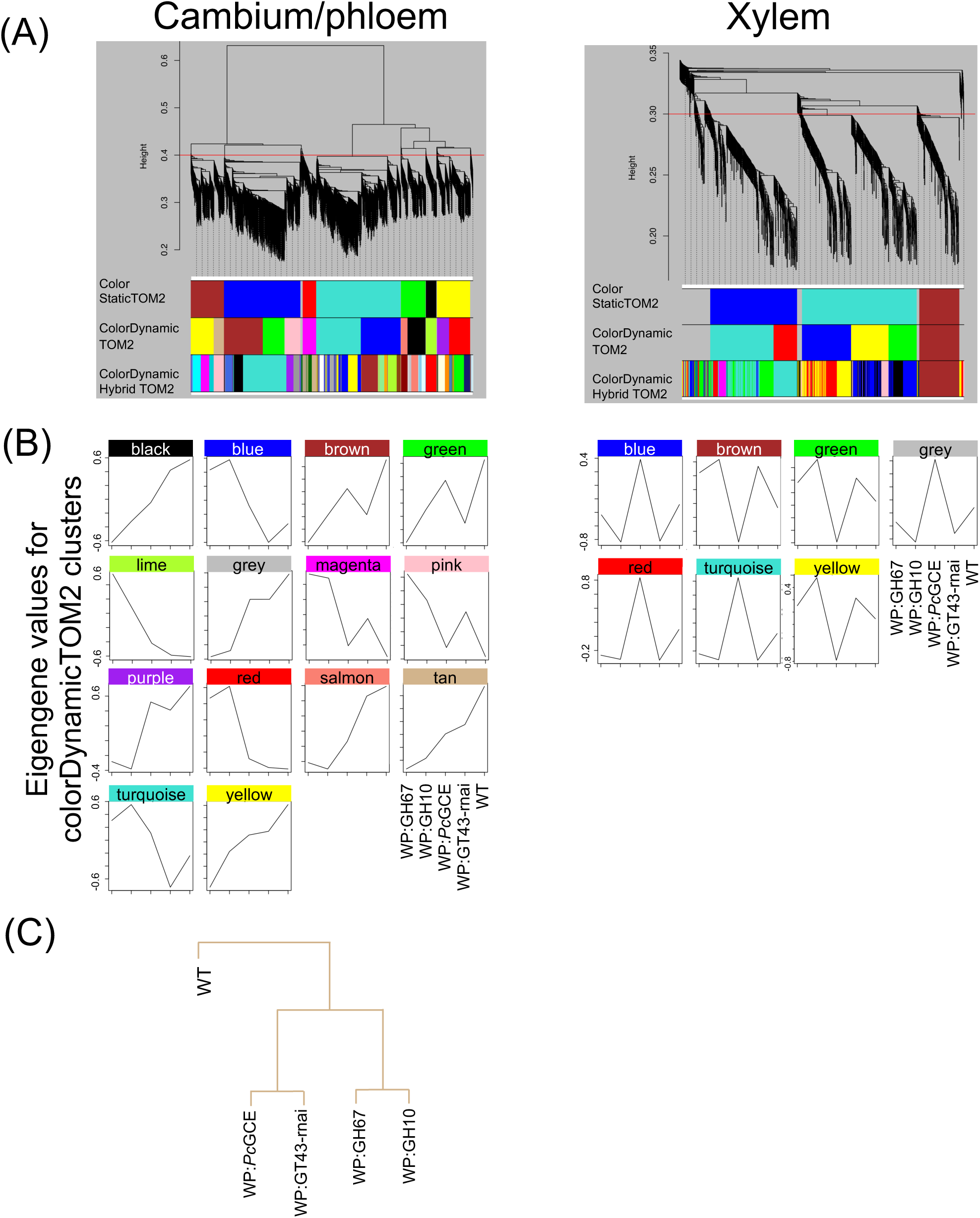
Clustering of genes based on correlations in different xylan-modified lines and among metabolomics, hormonomics, SilviScan and growth data. **(A)** Gene clusters, shown as dendograms for cambium/phloem and developing xylem tissues, obtained by weighted gene correlation network analysis (WGCNA). The color rows below the dendrograms indicate module membership identified by three different methods: the ColorStatic method (1^th^ band), the ColorDynamic method (2^nd^ band) and the ColorDynamicHybrid method (3^rd^ band). **(B)** The patterns of gene expression in the different lines for the clusters of the second band. **(C)** A dendrogram of the tan cluster showing the grouping of genotypes of transgenic lines, separately from the wild type (WT).

We further analyzed if the genes of the Tan cluster were coregulated during normal wood formation in aspen using the gene network analysis tool and AspWood dataset (Sundel et al., 2017) available at the PlantGenIE website (https://plantgenie.org/). This detected three gene networks, named Tan 1, Tan 2 and Tan 3, grouping 21, 22 and 88 genes, respectively (**Figure 8; Supplementary Table S9**). The Tan 1 network grouped genes that tended to be downregulated in transgenic lines but very few of them were significantly downregulated (P_adj_ ≤ 0.01, | log_2_(fold change) | ≥ 0.584). It was dominated by genes encoding mitochondrial proteins essential for respiration and ROS production. It also included homologs of *Arabidopsis* genes having function in ABA signaling, *RGLG1* (Wu et al., 2016) and *SCD2* (Hou and Shen, 2020) which was consistent with the decreased levels of ABA in the cambium/phloem tissues (**Figure 4**). Moreover, the Tan 1 network contained important developmental regulators, homolog of *VND1*, a NAC transcription factor triggering differentiation of tracheary elements and secondary wall formation (Ohashi-Itoh et al., 2024), *ARF3* regulating meristem size (Sang et al., 2022) and *ANGUSTIFOLIA (AN)* regulating polar cell growth (Chen et al., 2022) and homeostasis between SA and JA/Et levels (Xie et al., 2020). The Tan 2 network (**Figure 8; Supplementary Table S9**) grouped genes that tended to be upregulated in transgenic lines. It included homologs of regulators of vascular tissue development, *VPNB1* (Podia et al., 2018), *CLAVATA1* (*CLV1;* Zhang et al., 2024) and *ALTERED PHLOEM DEVELOPMENT* (*APL;* Bonke et al., 2003), and genes related to phloem function, like a homolog of *CALLOSE SYNTHASE 7* (*CalS7;* Xie et al., 2011), and two sucrose synthase-encoding genes, *PtSUS5* and *PtSUS6* (Kumar et al., 2019). I also contained other potential developmental regulators such as *NAC028* and *SQUAMOSA PROMOTER-BINDING PROTEIN LIKE4* (*SPL4*) as well as several genes encoding ankyrin repeat proteins. The Tan 3 network was the largest and it included genes having a weak tendency for downregulation in transgenic lines. Most of them were encoding ribosomal proteins and proteins related to translation (**Figure 8; Supplementary Table S9**). The remaining genes of this network were involved in the regulation of secondary growth, for example, regulation of cell division and meristem size, such as cyclin D (*CYCD3;3*; Dewitte et al., 2007), *PROHIBITIN3* (*PHB3*; Kong et al., 2018) and histone deacetylase (*HDA3/HDT1*; Zhang et al., 2019), and signaling such as homologs of *RACK1B* involved in ABA responses (Guo and Sun, 2017) and *ETO1* limiting ethylene biosynthesis by targeting a subset of ACC synthases to ubiquitination (Lee et al., 2021). Incidentally, three ubiquitin ligase-related genes were in this network, homologs of *PEX10* (Burkhart et al., 2014), *AIRP2* (Cho et al., 2014) and *AT1G30070* (**Figure 8; Supplementary Table S9**). The genes identified in these three networks constitute candidates for the fine tuning of gene expression in the cambial tissues in response to signals from adjacent xylem cells depositing secondary walls.

**Figure 8.**
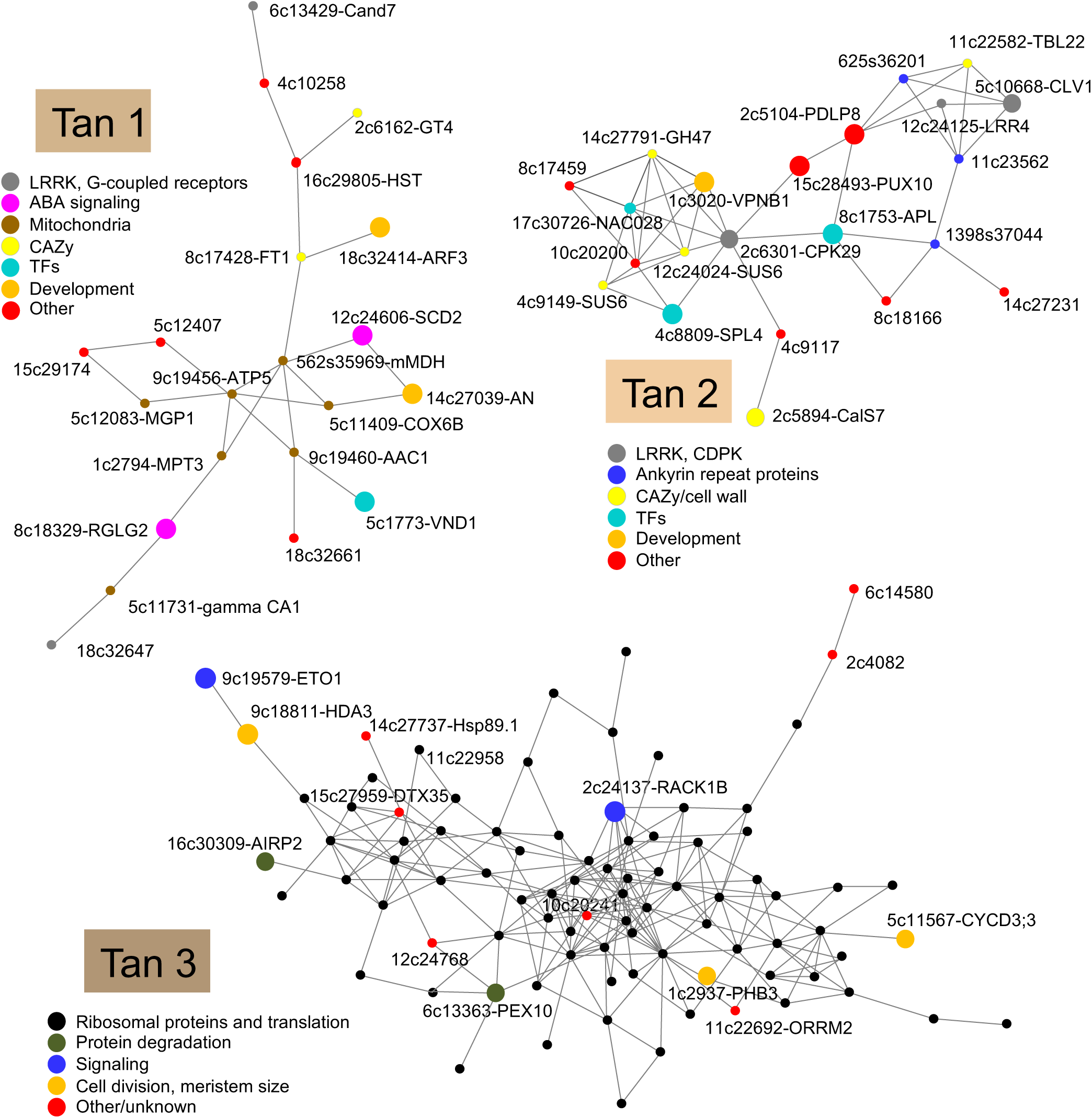
Co-expression networks of genes of WGCNA Tan cluster that were correlated with growth and were distinctively expressed in xylan-modified lines with increased growth compared to wild type across wood-forming aspen tissues. Data from AspWood (https://plantgenie.org/). Nodes are labeled with *P. tremula* gene ID that follows “Potra2n”and *Arabidopsis* gene name, if identified. In Tan3 network, the gene codes for ribosomal proteins encoding genes were omitted for clarity. The lists of genes for all networks are given in **Table S9**. Larger node symbols were used for genes discussed in the text.

## Discussion

Despite many papers reporting effects of secondary wall integrity impairments on various physiological responses, it is still unclear if there are common physiological responses and common signaling pathways that link the different secondary wall defects to specific outcomes. Here, we addressed this question by investigating the physiological and molecular consequences of different types of xylan alterations in the secondary walls in hybrid aspen.

Xylan is a key hemicellulose of secondary wall layers in angiosperms. Based on extensive analyses of *Arabidopsis* it is known that mutants defective in xylan biosynthetic genes affect growth to various degrees depending on redundancy, gene dosage and the type of xylan defect. Mutants defective in meGlcA side chains have no or much more attenuated effect on growth compared to mutants in xylan backbone or acetylation which can be dwarf (Wu et al., 2009; 2010; Lee et al., 2010; Mortimer et al., 2010; Ramírez and Pauly, 2019). Here, we compared effects of fungal hydrolases targeting either xylan backbone or meGlcA side chain, either ubiquitously or specifically in cells forming secondary walls in aspen. As in *Arabidopsis*, the observed effects on growth were variable, with xylanases having more detrimental effect than an α-glucuronidase, and severity of growth inhibition depending on xylanase expression levels (**Figure 1**). However, unlike in *Arabidopsis*, we observed that, in some instances, the growth of entire plants was stimulated by xylan impairment confirming previous observations (Biswal et al, 2015; Derba-Maceluch et al., 2015; Ratke et al., 2018). This occurred only when hydrolases or suppression of native xylan biosynthetic genes were targeted specifically to cells depositing secondary walls, and it concerned 30% of such lines, whereas in 95% of lines with ubiquitously targeted xylan modification the growth was inhibited. This suggests that primary and secondary wall integrity impairments can have opposite effects on growth. Indeed, the primary wall impairments reportedly inhibited growth (Hématy et al., 2007; Gigli-Bisceglia et al., 2018; Dünser et al., 2018; Sampathkumar et al., 2019). Inhibition of cell division is thought to be mediated by a pathway dependent on nitrate and cytokinin reductases controlling cytokinin levels (Gigli-Bisceglia et al., 2018), whereas inhibition of cell expansion is controlled by *THE1-* or *FER* – mediated pathways (Hématy et al., 2007; Dünser et al., 2018). To identify molecular players involved in growth stimulation induced by secondary wall impairment, we used hormonomics, metabolomics and transcriptomics analyses in the xylem cells depositing secondary walls, *i.e.* at the site of wall damage perception, and in adjacent cambial region tissues, *i.e.* where the growth is stimulated. The summary of our findings is presented in **Figure 9**.

**Figure 9.**
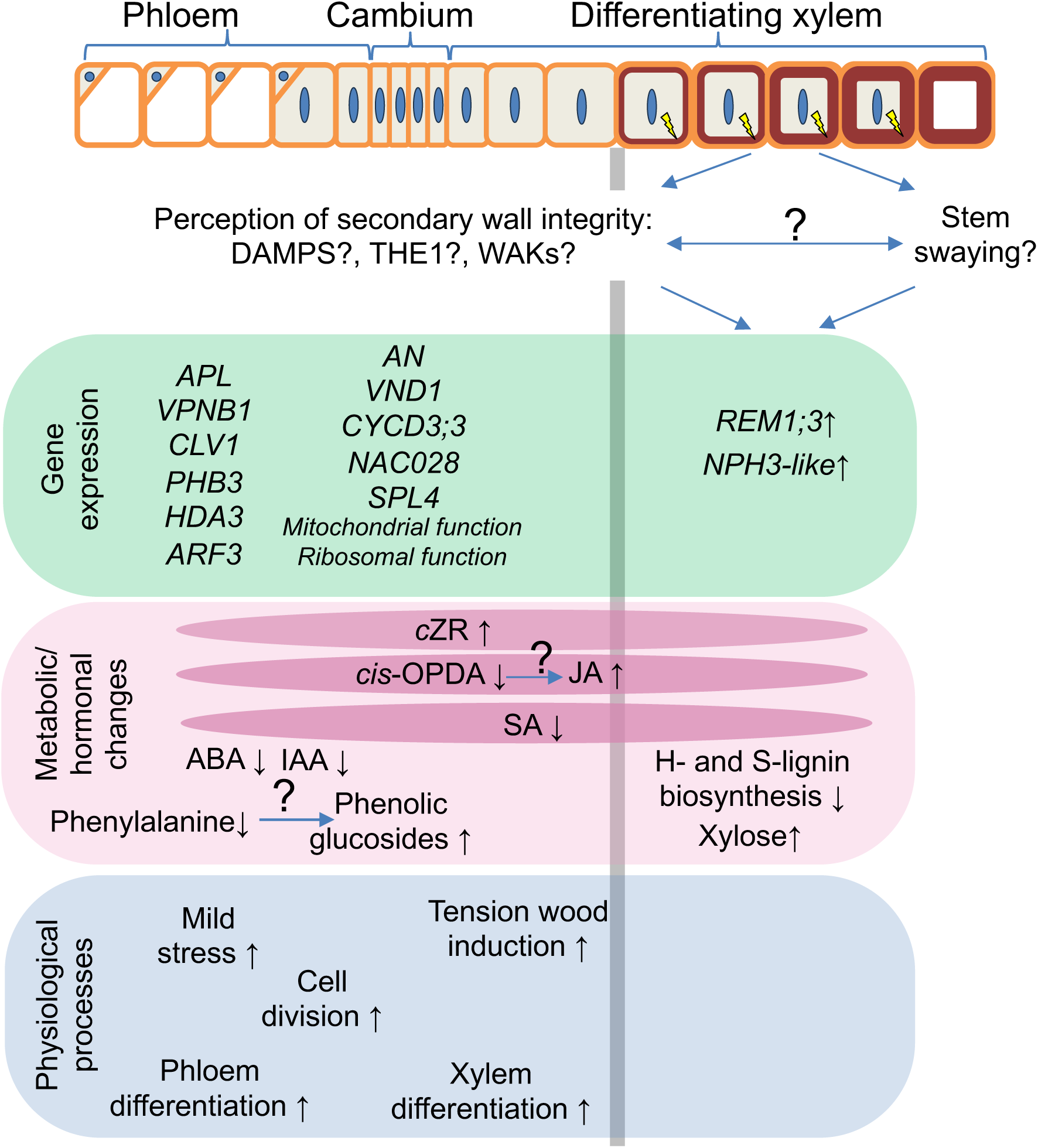
Diagram illustrating changes in the physiological processes, metabolic/hormonal status and gene expression which are induced in wood-forming tissues when secondary cell wall xylan is mildly modified. Increased stem swaying is expected when xylan integrity is impaired, but the relationship between stem swaying reaction and putative secondary cell wall defect perception is unknown. We suggest that DAMPs such as oligogalacturonides and/or xylobiose (Devangan et al., 2023; Sivan et al., 2025) are involved in the integrity perception along with receptors such as THE1 and WAKs. The subsequent signaling induces REM1;3 and NPH3-like genes in the xylem that could further regulate the cell wall integrity responses. Hormonal changes can occur in both cambium/phloem and developing xylem tissues (marked by darker pink ovals) include a conversion of *cis*-OPDA to JA, a decrease in SA, possibly resulting from decreased biosynthesis in the phloem, and an increase in *c*ZR that is probably transported from the phloem. ABA and IAA decrease in the cambium/phloem. Tension wood is induced in differentiating xylem. Farther in the cambial region tissues, cell division and xylem/phloem differentiation are activated. Phenylalanine might be converted into phenolic glycosides, indicating a possible stress response. Genes responding in the cambium/phloem are related to mitochondrial and ribosomal functions, cell cycle, meristem size regulation and vascular differentiation. Grey vertical line indicates the approximate split between cambium/phloem and differentiating xylem tissues that were analyzed.

Expression of three genes was affected at the cell wall damage site (i. e. in the secondary wall forming xylem) in all xylan-altered lines (**Figures 6, 9**). Of these, two, *REM1.3* encoding remorin and *NRL2* of *NPH3* family were upregulated in transgenic lines and they are of particular interest. Remorins are essential for lipid raft organization and symplasmic signaling. In *Arabidopsis*, REM1.3 protein is rapidly phosphorylated by the CALCIUM-DEPENDENT PROTEIN KINASE 3 (CDPK3) (Perraki et al., 2018) in response to oligogalacturonides (Kohorn et al., 2016b), which regulates nanodomain organization and restricts plasmodesmatal transport downstream SA signaling (Huang et al., 2019). Decreased SA concentration observed in transgenic lines (**Figures 4, 9**) would then increase symplasmic transport. Regulation of symplasmic transport is relevant in case of secondary wall integrity signaling because cell wall damage perception must occur specifically during the secondary wall layer deposition *i.e.* when cell growth is already finished, and its perception needs to induce a mobile signal transmitted to the dividing and growing cells, and possibly spreading to other organs to affect growth. Another gene upregulated in all studied xylan-modified lines was a homolog of *NRL2* from the *NPH3* family which encodes proteins interacting with blue-light receptor-like kinases called phototropins and mediating blue-light phototropic responses (Christie et al., 2018). NRL2 protein was found to interact with PHOTOTROPIN 1 (PHO1) but the interaction was not affected by light (Sullivan et al., 2009), suggesting a role in a different process than light signaling. Therefore both, *REM1.3* and *NRL2,* are key candidates for secondary wall integrity sensing and signaling which leads to better growth, and should be further analyzed.

Many changes in metabolites and hormones were observed in common between the cambium/phloem and xylem samples (**Figures 4, 9**). This could be related to increased symplasmic transport, as discussed above or to increased apoplasmic transport due to cell wall defect. Among the most notable changes in hormonal status in lines with impaired xylan there was a tendency for increase in JA and decrease in its precursor, *cis-*OPDA, which could indicate increased biosynthesis of active jasmonates in the developing xylem (**Figure 4**). This increase could play a role in growth stimulation observed in the transgenic lines as JA is known for its stimulatory effect on cambial divisions and xylem formation (Sehr et al., 2010; Jang et al., 2017). Decreased ABA levels in the transgenic lines (**Figure 4**) could also play an important role in secondary wall integrity signaling. ABA is known for its growth inhibitory effects (Popko et al., 2010). Strongly decreased ABA levels accompanied enhanced cambial activity and phloem production in lines with severe secondary cell wall damage (Sivan et al., 2025). Recently, cell wall damage induced by isoxaben was observed in the central part of the *Arabidopsis* root, corresponding to the stele, where it was shown to induce THE1 that mediated JA signaling (Bacete et al., 2022). The treatment also reduced ABA in adjacent tissues, also in a THE1-dependent manner. These observations along with the data presented in this paper strongly implicate reduced ABA and increased JA signaling downstream THE1 as a common response to secondary cell wall defects.

Interestingly, similar changes in the hormones (decreased *cis*-OPDA and reduced ABA concentrations) were observed in wood-forming tissues of stems subjected to stem swaying (Urbancsok et al., 2023). This treatment also stimulated plant growth and xylem production. Moreover, it induced increased formation of tension wood, similar to effects of xylan impairment (**Figures 2, 9**). These similarities indicate that pathways between mechanical stress perception during stem swaying and secondary cell wall damage overlap and possibly the changes reported here in the xylan-modified plants are mediated *via* stem swaying. It would be therefore crucial to determine if the stem swaying is needed for growth stimulation in secondary wall-integrity compromised plants. Secondary wall damage perception likely also involves DAMPs such as xylobiose (Devangan et al., 2023), as found in the case of xylanase-overexpressing aspen (Sivan et al., 2025). Here we found that aspen with xylan defects and increased growth showed increases in xylose and derived sugars (**Supplementary Table S4**) which could originate from xylobiose or xylan. Oligogalacturonides could also play a role as DAMPs when xylan integrity is altered, because xylan and pectin networks are likely interconnected in the wood (Biswal et al., 2014).

At the site of increased growth, i.e. in the cambial region tissues, transcriptomic changes were subtle, but the WGCNA clustering identified the Tan cluster of co-regulated transcripts in transgenic lines which correlated with growth promoting hormones, like IAA and tZ, and metabolites related to amino-acids and phenylpropanoids (**Figure 7, Supplementary Tables S6, S8**). These results support common genetic machinery responsible for growth stimulation. Interestingly, these genes formed co-expression networks during normal process of xylogenesis (**Figure 8, Supplementary Table S9**), which could operate during the monitoring of secondary wall integrity and fine-tuning of xylogenesis with growth. This fine-tuning appears to specifically regulate different groups of genes. One of them represents phloem biogenesis-related genes illustrated by the Tan 2 network, grouping genes related to vascular development, phloem fate specification and phloem function. Incidentally, our previous analyses showed increased phloem production when xylan structure was heavily affected (Sivan et al., 2025). Another group represents genes regulating mitochondrial function as exemplified by the Tan 1 (**Figure 8, Supplementary Table S9**). This is particularly intriguing given that the defects in mitochondrial functions were shown to confer the resistance to cell wall damage caused by inhibition of cellulose biosynthesis (Hu et al., 2016).

Among the most and commonly increased metabolites in transgenic lines there were phenyl glycosides (**Figures 5, 9, Supplementary Table S4**) which are known to accompany biotic stress responses (Babst et al., 2010). Thus, the modification of xylan could results in activation of biotic stress responses, as also known for other cases of secondary wall impairment (Hernández-Blanco et al., 2007; Gallego-Girlado et al., 2011a, Pogorelko et al., 2013; Pawar et al., 2016; Molina et al., 2021).

In conclusion, we demonstrated that common physiological, hormonal, metabolic and transcriptional responses are activated in wood-forming tissues when xylan is impaired in developing secondary walls (**Figure 9)**. How cell wall defects are perceived in the xylem cells is still unknown but we have identified two genes, *REM1;3* and *NRL2*, which are induced in the xylem upon secondary wall xylan damage and could regulate downstream responses to impaired secondary walls. These responses are partially non-cell autonomous involving changes in the cambial meristem and further affecting growth of entire plants. The identified genes and hormones likely coordinate the process of secondary wall formation with cambial activity and vascular differentiation during secondary growth.

## Short legends for supplementary data

**Figure S1.** Analysis of transcriptomic changes in the cambium/phloem tissues of transgenic lines compared to wild type using WGNCA.

**Figure S2.** Analysis of transcriptomic changes in the xylem tissue of transgenic lines compared to wild type using WGNCA.

**Table S1**. Primers used in this study.

**Table S2**. Levels of all compounds detected by hormonomics analysis in developing wood tissues of the selected transgenic lines with modified xylan and increased growth.

**Table S3**. SilviScan wood traits that were not affected in the selected transgenic lines with modified xylan and increased growth

**Table S4**. All compounds detected by metabolomics analysis in developing wood tissues of the transgenic lines with modified xylan and increased growth.

**Table S5**. Genes differentially expressed (|Log_2_ (fold change) |≥ 0.584 and Padj≤0.01) in wood forming tissues of at least one transgenic line compared to wild type.

**Table S6**. Clusters of gene co-regulated in the cambium/developing phloem of transgenic lines and wild type according to WGCNA analysis.

**Table S7**. Clusters of gene co-regulated in the xylem of transgenic lines and wild type according to WGCNA analysis.

**Table S8**. WGCNA clusters of co-regulated genes correlated with different variables corresponding to growth, hormones, and metabolites.

**Table S9**. Co-expression networks in the wood-forming aspen tissues for the Tan WGCNA cluster of co-regulated genes correlated with different variables including growth, hormones, and metabolites.

## Acknowledgement

We acknowledge the Umeå Plant Science Centre (UPSC) Tree Phenotyping Platform, Bioinformatics Facility, Microscopy Facility, Biopolymer Analytical Platform and the Swedish Metabolomics Centre in Umeå (www.swedishmetabolomicscentre.se). We thank Thomas Grahn (RISE) for help with NIR-estimations of tension wood.

## Author Contributions

JU supervised the experiments, processed wood material, extracted RNA, analyzed transgene expression and prepared samples for omics analyses. END analyzed roots, designed and performed bioinformatic analyses and prepared figures. MDM created transgenic aspen and collected samples for SilviScan analysis. PS performed wood chemistry analyses. JU, END and FRB carried the greenhouse experiment and tree phenotyping. MM analyzed metabolomics data and prepared them for publication. ZY and GS carried out wood SilviScan and NIR analyses. JŠ, KC and MK analyzed hormones and related compounds. EJM designed and coordinated the research, secured the funding, and wrote the paper with contributions from all authors.

## Data availability

The raw RNA-Seq data that support the findings of this study are available in the European Nucleotide Archive (ENA) at EMBL-EBI (https://www.ebi.ac.uk/ena/browser/home), under accession ID PRJEB82792.

